# Does closing the eyes enhance auditory attention? Eye closure increases attentional alpha-power modulation but not listening performance

**DOI:** 10.1101/455675

**Authors:** Malte Wöstmann, Lea-Maria Schmitt, Jonas Obleser

**Author notes:** Author correspondence: Malte Wöstmann & Jonas Obleser, Maria-Goeppert Straße 9a, 23562 Lübeck, Germany. M.W. and L.-M.S. contributed equally and share first authorship.

## Abstract

In challenging listening conditions, closing the eyes is a strategy with intuitive appeal to improve auditory attention and perception. On the neural level, closing the eyes increases the power of alpha oscillations (∼10 Hz), which are a prime signature of auditory attention. Here, we test whether eye closure benefits neural and behavioural signatures of auditory attention and perception. Participants (*N* = 22) attended to one of two alternating streams of spoken numbers with open or closed eyes in a darkened chamber. After each trial, participants indicated whether probes had been among the to-be-attended or to-be-ignored numbers. In the electroencephalogram, states of relative high versus low alpha power accompanied the presentation of attended versus ignored numbers. Importantly, eye closure did not only increase the overall level of absolute alpha power but also the attentional modulation thereof. Behaviourally, however, neither perceptual sensitivity nor response criterion was affected by eye closure. To further examine whether this behavioural null-result would conceptually replicate in a simple auditory detection task, a follow-up experiment was conducted that required participants (*N* = 19) to detect a near-threshold target tone in noise. As in the main experiment, our results provide evidence for the absence of any difference in perceptual sensitivity and criterion for open versus closed eyes. In sum, we demonstrate here that the modulation of the human alpha rhythm by auditory attention is increased when participants close their eyes. However, our results speak against the widely held belief that eye closure per se improves listening behaviour.

---

When we listen to faint sounds in noise-contaminated environments, we sometimes close our eyes with the intention to strengthen the focus on auditory sensory input. In theory, eye closure has been proposed to free perceptual and cognitive resources (Glenberg, Schroeder, & Robertson, 1998; Vredeveldt, Hitch, & Baddeley, 2011) to focus attention on non-visual sensory information. But does eye closure indeed have the potency to enhance auditory attention and perception?

There is some evidence that closing the eyes can improve performance in certain sensory and cognitive tasks. Somatosensory perception thresholds decrease when participants have their eyes closed compared to open, in an illuminated but even in a dark room (Brodoehl, Klingner, Stieglitz, & Witte, 2015; Brodoehl, Klingner, & Witte, 2015). This speaks to the general potency of eye closure to enhance perceptual processing. Evidence for enhanced cognitive capacity through eye closure comes from studies showing that eyewitnesses’ recall of the details of a crime scene improves when they close their eyes during memory recall (e.g., Perfect et al., 2008; Vredeveldt, Baddeley, & Hitch, 2012; Vredeveldt et al., 2015). However, systematic investigations of the impact of eye closure on perceptual and cognitive functions across sensory modalities are sparse at best.

Evidence for listeners’ use of eye closure as a strategy to improve listening is mostly anecdotal. In his seminal paper establishing the Cocktail party effect, Colin Cherry (1953) studied listeners’ separation of two concurrent spoken messages and noted that “At the subjective level the subject reported great difficulty in accomplishing his task. He would shut his eyes to assist concentration.”^1^ Nowadays, it is common practice in auditory research to put efforts into avoiding that participants close their eyes in the laboratory. It is all the more surprising that there is to our knowledge no published work on the behavioural or neural consequences of closing the eyes during attentive listening (but see Götz et al., 2017 for initial evidence from a combined somatosensory and auditory study).

Indirect evidence in favour of an eye–closure-induced modulation of auditory processing comes from studies showing that eye closure in darkness increases the blood oxygen level dependent (BOLD) contrast in auditory cortex regions (Marx et al., 2004; Marx et al., 2003) and increases the acoustic reflex (Corcoran, Cleaver, & Stephens, 1980), that is, a sound-induced muscle contraction in the middle ear. Furthermore, eye position has been shown to impact activity in inferior colliculus (Groh, Trause, Underhill, Clark, & Inati, 2001) and auditory cortex (Werner-Reiss, Kelly, Trause, Underhill, & Groh, 2003) in primates. The present study aims at testing whether eye closure affects one of the arguably most abundant listening situations in everyday life, that is, listening to an auditory signal despite distraction.

Neuroscience has actually long established an indirect link between eye closure and auditory attention: It is known since the work by Hans Berger and Lord Adrian in the 1940s that eye closure and auditory attention induce a similar electrophysiological response: an increase in the power of alpha oscillations (∼10 Hz) at parietal and occipital scalp sites (Adrian, 1944; Adrian & Matthews, 1934; Berger, 1929). More recently, high alpha power has been interpreted as a means to relatively inhibit neural processing in task-irrelevant brain areas (Foxe & Snyder, 2011; Jensen & Mazaheri, 2010). Among other cognitive processes that modulate the power of alpha oscillations, auditory tasks increase alpha power in parieto-occipital cortex regions, possibly in order to increase attention to the auditory modality (e.g., Fu et al., 2001; Strauß, Wöstmann, & Obleser, 2014; Weisz, Hartmann, Müller, Lorenz, & Obleser, 2011).

Alpha power modulations do covary with behavioural performance in auditory attention and memory paradigms (Wilsch, Henry, Herrmann, Maess, & Obleser, 2015; Wöstmann, Herrmann, Maess, & Obleser, 2016; Wöstmann, Herrmann, Wilsch, & Obleser, 2015). Furthermore, we have recently shown that modulatory brain stimulation at alpha frequency influences the degree of distraction by irrelevant speech (Wöstmann, Vosskuhl, Obleser, & Herrmann, 2018). It is thus likely that alpha power is of functional relevance to auditory attention. The rationale of the present study is the following: If the eye–closure-enhanced alpha oscillators in parieto-occipital cortex regions (partly) overlap with the alpha oscillators active during auditory attention, then closing the eyes should increase the auditory attention-induced modulation of alpha power. Beyond effects on neural oscillatory dynamics, we here test whether eye closure also has the potency to improve behavioural performance in auditory attention and perception.

The primary goal of this study was to test whether auditory attention and eye closure interactively modulate alpha power and listening behaviour. To this end, we utilised an auditory temporal attention task that is known to induce attentional modulation of alpha power in similar brain areas as eye closure, that is, in parieto-occipital cortical regions. In this respect, spatial attention tasks or tasks involving continuous attention to one of several simultaneous auditory streams would not have been well suited paradigms. Spatial attention tasks induce alpha power changes between the two hemispheres (as opposed to bilateral, parieto-occipital regions; e.g., Wöstmann et al., 2016) and tasks involving continuous attention to one of several simultaneous streams primarily evoke selective phase-locking of low-frequency oscillations but no modulation of alpha power (e.g., Fiedler, Wöstmann, Herbst, & Obleser, 2019). Although the auditory evoked response is another neural signature modulated by attention, no modulation of the evoked response by eye closure has been reported to our knowledge. We did not expect interactive effects of attention and eye closure on the auditory evoked response, which was thus no outcome measure of primary interest in the present study.

In the main experiment, participants (*N* = 22) selectively attended to one of two alternating streams of spoken numbers. Auditory attention was operationalized neurally as the difference in alpha power for attended versus ignored numbers, and behaviourally as accuracy in classifying probe numbers as attended versus ignored. To rule out that effects of eye closure were confounded by participants perceiving visual input (versus darkness), participants performed the task in a darkened chamber, with eyes open or closed during half of the blocks of the experiment. In a follow-up experiment, we tested the impact of eye closure on the detection of non-speech stimuli in noise in a separate sample of participants (*N* = 19).

## Materials and Methods

### Participants

Healthy, right-handed, native German-speaking participants took part in the main experiment (*N* = 22; 12 females; age range 19–31 years, *M* = 24.68, *SD* = 3.06) and in the follow-up experiment (*N* = 19; 17 females; age range 20–31 years, *M* = 23.89, *SD* = 4.04). In the main experiment, the data of one additional participant were excluded from all analyses due to noise-contaminated EEG signals. All participants signed informed consent and were financially compensated for participation. All procedures were approved by the local ethics committee of the University of Lübeck.

### Auditory materials in the main experiment

Speech stimuli consisted of two sets of 72 spoken numbers from 21 to 99 excluding integer multiples of 10. One set was spoken by a trained German female speaker (used in previous studies; Wöstmann, Herrmann, et al., 2015; Wöstmann, Schröger, & Obleser, 2015) and the other by a male speaker. For instructions, a third set of numbers and spoken instructions was recorded from a different German female speaker. Praat software (version 6.0.18) was used to shift the formant frequencies of this third speaker slightly towards a male voice (formant shift ratio of 1.3), in order to render the instructor voice gender-neutral. Speech recordings were made in a soundproof room and digitized at a sampling rate of 44.1 kHz. Numbers ranged in duration from 1.01 to 1.26 s (*M* = 1.12, *SD* = 0.06) and comprised four syllables.

For each trial, five randomly selected female-voiced numbers and five randomly selected male-voiced numbers were alternately assigned within the ten-number sequence. Half of the trials started with a female-voiced number, that is, all odd number positions were occupied by female-voiced numbers and all even number positions by male-voiced numbers (and vice versa for trials starting with a male-voiced number). All ten numbers in a trial were different from and numerally non-successive to one another. The time interval between the onsets of two sequent numbers was 1.33 s. Number sequences were energetically masked by continuous white noise (signal-to-noise-ratio of +10 dB) with onset and offset raised cosine ramps of 0.15 s preceding and ensuing a number sequence, respectively. Number sequences in background noise ranged in duration from 13.28 to 13.53 s (*M* = 13.39, *SD* = 0.06) depending on the duration of the last number.

### Procedure of the main experiment

The experiment was designed in a way that it could be performed with open but also with closed eyes. Each trial started with the spoken instruction to attend to the female- or male-voiced numbers. After 0.9 s (randomly jittered 0.6–1.2 s), a sequence of ten numbers was presented. Another 0.9 s (randomly jittered 0.6–1.2 s) after number sequence offset, three probe numbers of this trial were uttered by the instructor voice. After each probe, participants responded via button press on a computer mouse whether the probe was one of the to-be attended numbers (left button, ‘Yes, attended’-response) or not (right button, ‘No, ignored’-response). There was no time limit for a response (Fig. 1A).

**Figure 1.**
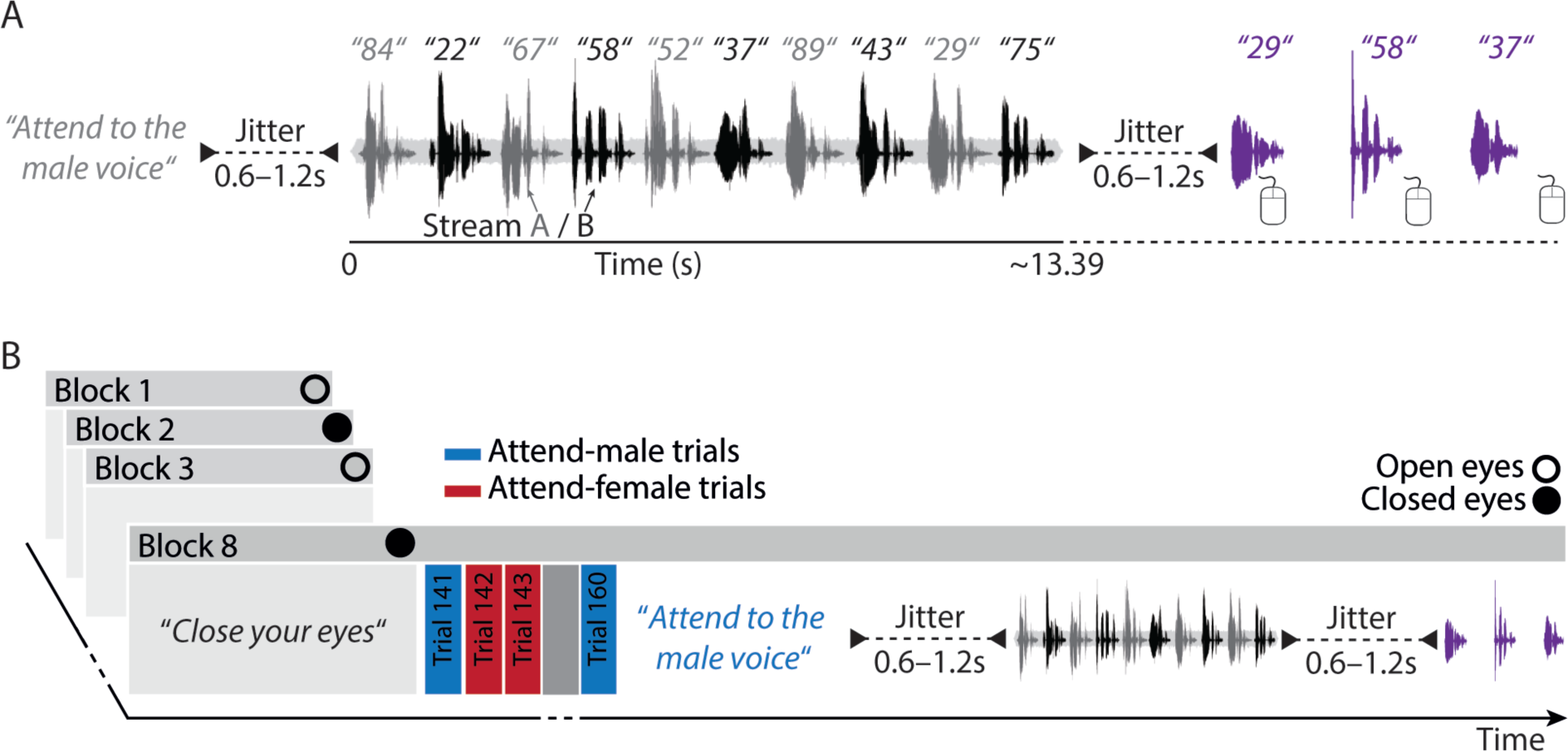
Task design of the main experiment. (A) In each trial, listeners attended to speech stream A (grey numbers) or B (black numbers), each composed of five spoken German numbers embedded in a white noise masker (+10 dB SNR). At the end of each trial, three probe numbers were presented (purple). Participants had the task to indicate with a mouse button press, whether the probed number was ‘attended’ or ‘ignored’. (B) Each one of eight blocks started with the instruction to keep the eyes open or closed during the entire block. For each trial within a block, the task instruction to attend to the male or female voice was randomly assigned.

Numbers from the temporally middle positions were probed more frequently. In detail, for each condition in the 2 (Eyes: open vs. closed) × 2 (Attended voice: male vs. female) × 2 (Attended stream: A, odd positions vs. B, even position) design, 60 to-be-probed number positions (for 20 trials with three probes each) were predefined: Number positions 1, 2, 9, and 10 were probed three times each; number positions 3, 4, 7, and 8 were probed seven times each; and number positions 5 and 6 were probed ten times each.

During the whole trial a grey fixation circle was presented in the middle of a black computer screen placed in front of participants. Trials were separated by a time interval of 1.3 s. All ten numbers presented in a trial were excluded from reoccurring in the subsequent trial.

Twenty trials of the experiment formed a block; a total of 8 blocks resulted in 160 trials per participant. At the beginning of a block participants were presented a spoken instruction to either close their eyes or to keep them open. Blocks were alternately worked through with open and closed eyes, whereby half of the participants completed the first block with open eyes. The entire experiment was performed in a darkened chamber. The light was turned on in between blocks for a self-paced break of at least one minute.

Stimulation was controlled by Presentation software (version 18.3; Neurobehavioural Systems). Auditory stimuli were presented diotically (i.e., to both ears and spatially non-separated) via headphones (Sennheiser HD 25-1 II). During the whole procedure participants were monitored via infrared camera to ensure they consistently kept their eyes open or closed in agreement with the instruction. The experiment took approximately 75 minutes to complete (i.e., ∼8 minutes per block).

### Auditory materials in the follow-up experiment

The target tone was a 1000-Hz sine tone of 100-ms duration and 10-ms linear onset and offset ramps. The distracting background noise was bandpass-filtered random noise (500–1500 Hz) of 3-s duration and 50-ms linear onset and offset ramps. For each trial of the follow-up experiment, a different background noise stimulus was created.

### Procedure of the follow-up experiment

As the main experiment, the follow-up experiment was designed in a way that it could be performed with open but also with closed eyes. Participants had the task to report detection of the target tone in noise using a computer mouse (left button: ‘Yes’, tone detected; right button: ‘No’, no tone detected). The target tone was present on 50% of trials. For target-present trials, the onset of the target tone was placed at a random millisecond-value between 1500 and 2100 ms following background noise onset. The experiment contained 600 trials, divided in 12 blocks à 50 trials (25 target-present trials and 25 target-absent trials). Every other block was performed with closed versus open eyes, with participants with odd versus even experiment-IDs starting with open versus closed eyes, respectively. Pre-recorded instructions in the beginning of each block (“Keep your eyes open/closed”) and in the end of each block (“This block is finished, take a brief break”) were spoken by a male voice. All auditory stimuli were presented diotically via headphones (Sennheiser HD 25-1 II). The follow-up experiment took approximately 75 minutes to complete. The experiment was performed in a darkened chamber, with the computer monitor switched off for the entire duration of the experiment. As in the main experiment, participants were monitored via infrared camera to ensure they consistently kept their eyes open or closed in agreement with the instruction. The light was switched on in the breaks in-between blocks of the experiment.

Prior to the start of the follow-up experiment, we titrated the Signal-to-Noise Ratio (SNR) of the target tone in noise for each individual participant with eyes open and the light switched on. To this end, participants performed a one-up one-down staircase procedure to determine the SNR for a proportion correct of 0.5 for target-present trials. Note that a proportion correct of 0.5 for target-present trials is typically accompanied by a somewhat higher proportion correct for target-absent trials (see Fig. 5E). Subsequently, participants performed a training block including 50 trials (25 target-present trials). The main experiment started if proportion correct averaged across target-present and target-absent trials was close to 0.65 (between 0.55 and 0.75). Otherwise, the SNR was adjusted and the training block was repeated.

**Figure 5.**
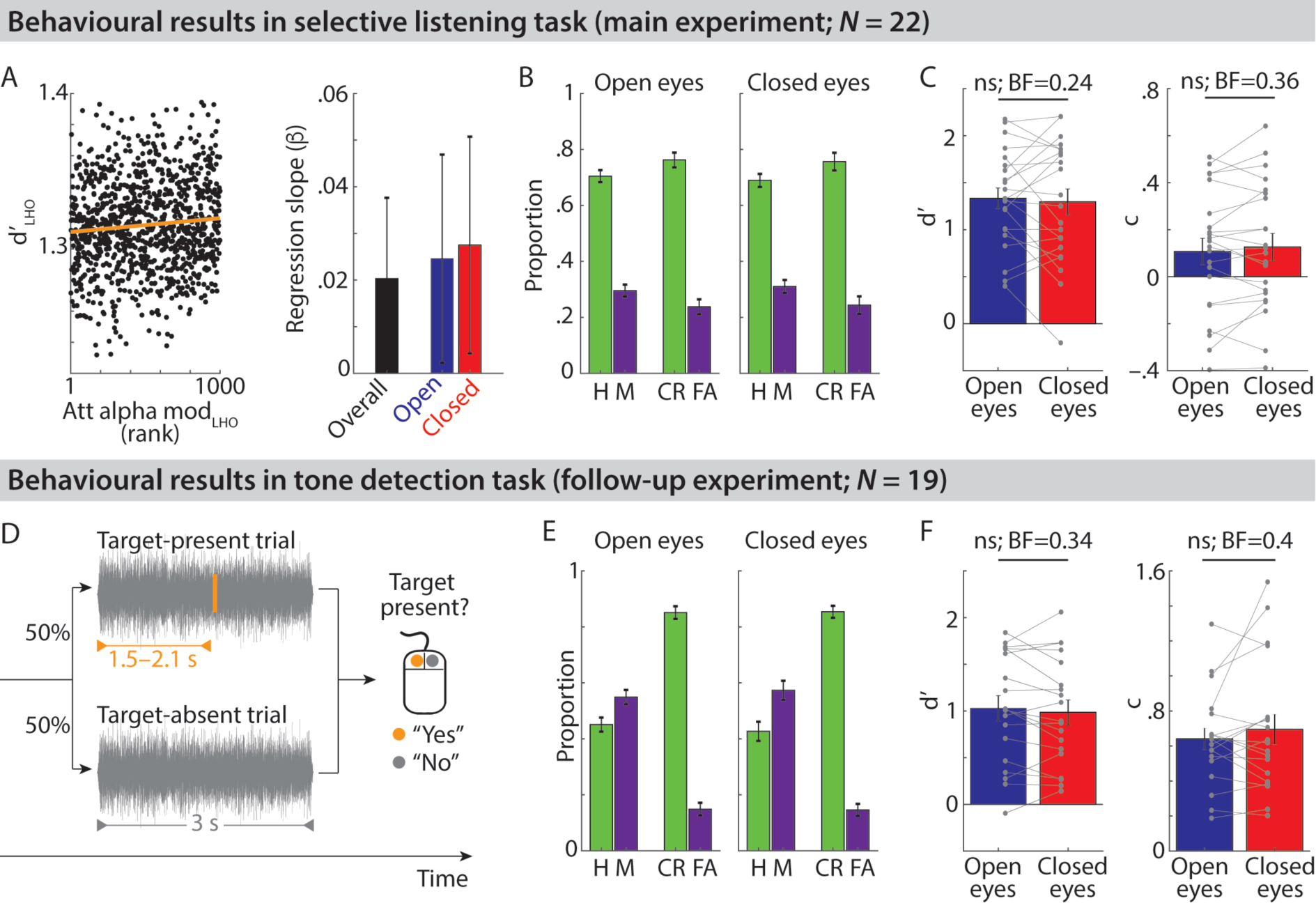
Modulation of behavioural performance in main- and follow-up experiment. (A) Scatterplot shows sensitivity (d’_LHO_) derived from 1,000 leave-half-out (LHO) subsamples of each participant’s data as a function of the attentional alpha modulation_LHO,_ averaged across *N* = 22 participants across eyes open and closed conditions in the main experiment. Orange line shows the least-squares linear fit. Bars show average slopes obtained through regression of d’_LHO_ on attentional alpha modulation_LHO_. (B) Bars show average proportions of response types in the main experiment, separately for open and closed eyes (H = Hit, M = Miss, CR = Correct Rejection, FA = False Alarm). (C) Bars and thin lines show respective average and single-subject sensitivity (d’) and criterion (c). (D) Task design of the follow-up experiment. Participants (*N* = 19) had to detect a tone (orange; which was present in 50% of trials) in noise (grey). (E & F) Same as A & B, but for the follow-up experiment. Error bars show ± 1 between-subject SEM. BF indicates the Bayes Factor for non-significant (ns) effects.

### Behavioural data analysis

In the main experiment, in order to base behavioural and EEG analyses on the same data, we only considered trials that passed the artefact rejection of EEG data (see below) for the behavioural data analysis.

On each trial, participants made three behavioural responses to judge the probes either as ‘Yes, attended’ or ‘No, ignored’. We calculated the proportions of Hits (correct ‘Yes, attended’-responses for to-be-attended probes), Misses (incorrect ‘No, ignored’-responses for to-be-attended probes), Correct rejections (correct ‘No, ignored’-responses for to-be-ignored probes), and False alarms (incorrect ‘Yes, attended’-responses for to-be-ignored probes). Sensitivity (d’) and response bias (c) were calculated according to signal detection theory (Macmillan & Creelman, 2005), separately for eyes open versus closed conditions. Larger values of d’ indicate higher sensitivity in internally separating attended from ignored numbers. Larger positive values of c indicate more conservative bias to respond ‘No, ignored’ more often.

In the follow-up experiment, we calculated the proportions of Hits (correct ‘Yes’-responses in target-present trials), Misses (incorrect ‘No’-responses in target-present trials), Correct rejections (correct ‘No’-responses in target-absent trials), and False alarms (incorrect ‘Yes’-responses in target-absent trials), separately for trials with open versus closed eyes. As for the main experiment, proportions of Hits and False alarms were used to obtain sensitivity (d’) and response bias (c).

### EEG recording and preprocessing

In the main experiment, participants were seated in an electrically shielded sound-attenuated EEG chamber. While they were prepared for EEG recordings, they performed a feedback-based training session (10 trials) with eyes open and lights on to get used to the task.

The electroencephalogram (EEG) was recorded at a sampling rate of 1 kHz with a pass-band from direct current (DC) to 280 Hz (actiCHamp, Brain Products). Sixty-four Ag/AgCl electrodes were fixed to an elastic cap (actiCAP) according to the extended 10-20 standard system at the following positions: FP1/2, AF3/4/7/8, F1–8, FC1–6, FT7–10, C1–6, T7/8, CP1– 6, TP7–10, P1–8, PO3/4/7/8, O1/2, FPz, AFz, Fz, FCz, Cz, CPz, Pz, POz, and Oz. The ground electrode was mounted on the forehead (FPz). During recording, electrodes were referenced against the left mastoid (TP9) and all impedances were kept below 5 kΩ.

EEG data were preprocessed and analysed using Matlab (version R2013b, MathWorks) and the FieldTrip toolbox (version 2012-12-16; Oostenveld, Fries, Maris, & Schoffelen, 2011). From the continuous data, epochs were extracted time-locked to the onset of acoustic stimulation in each trial (–1 to 14 s). Epochs were high-pass filtered at 1 Hz and low-pass filtered at 100 Hz using 6^th^ order Butterworth filters. Due to technical problems during recording, a small number of epochs (i.e., 1, 4, and 20 epochs) were missing for three participants, respectively.

For three participants, signals of bad EEG electrodes were interpolated. A bad electrode was defined as an electrode whose impedance was >5 kΩ at the end of the preparation for EEG measurements. The signal at electrode AF (one participant) and F6 (two participants) was replaced by a weighted average signal of neighbouring electrodes using the nearest-neighbours method.

An independent component analysis (ICA) was performed on each participant’s entire dataset (including eyes open and eyes closed conditions). Components’ time courses, topographies, and frequency spectra were inspected for artefacts (i.e., eye blinks, saccades, cardiac activity, and other muscle activity). On average, 50.48% of components (*SD* = 11.22%) were removed from the data. Further, single epochs were removed from the data if any channel’s activity range was greater than 250 μV within the time interval of the number stream in noise (0 to 13.53 s). On average 1.8% of epochs (*SD* = 2.64%) were excluded from further analyses. All epochs were re-referenced to the average of all electrodes.

In the follow-up experiment, the EEG was recorded but not analysed for the purpose of the present study.

### Analysis of oscillatory power

We performed two spectral analyses. First, non-time-resolved absolute oscillatory power across the average trial duration (0–13.39 s) was obtained using Fast Fourier Transform (FFT) with multitapering for frequencies 0.5–20 Hz in steps of 0.5 Hz with 1 Hz spectral smoothing, separately for trials with open versus closed eyes (Fig. 2A). Source localization of absolute alpha power (8–12 Hz) was performed using the Dynamic Imaging of Coherent Sources (DICS) beamformer approach (Gross et al., 2001). A standard headmodel (Boundary Element Method, BEM; 3-shell) was used to calculate leadfields for a grid of 1 cm resolution. For each participant, we calculated an adaptive spatial filter from the leadfield and the cross-spectral density of Fourier transforms of all trials (eyes open and closed) centered at 10 Hz with ±2 Hz spectral smoothing (resulting in a frequency range of 8–12 Hz) in the time interval 0–13.39 s relative to sound onset. This filter was applied separately to single-trial Fourier transforms of eyes open and closed conditions to obtain alpha power at each grip point (Fig. 2B).

**Figure 2.**
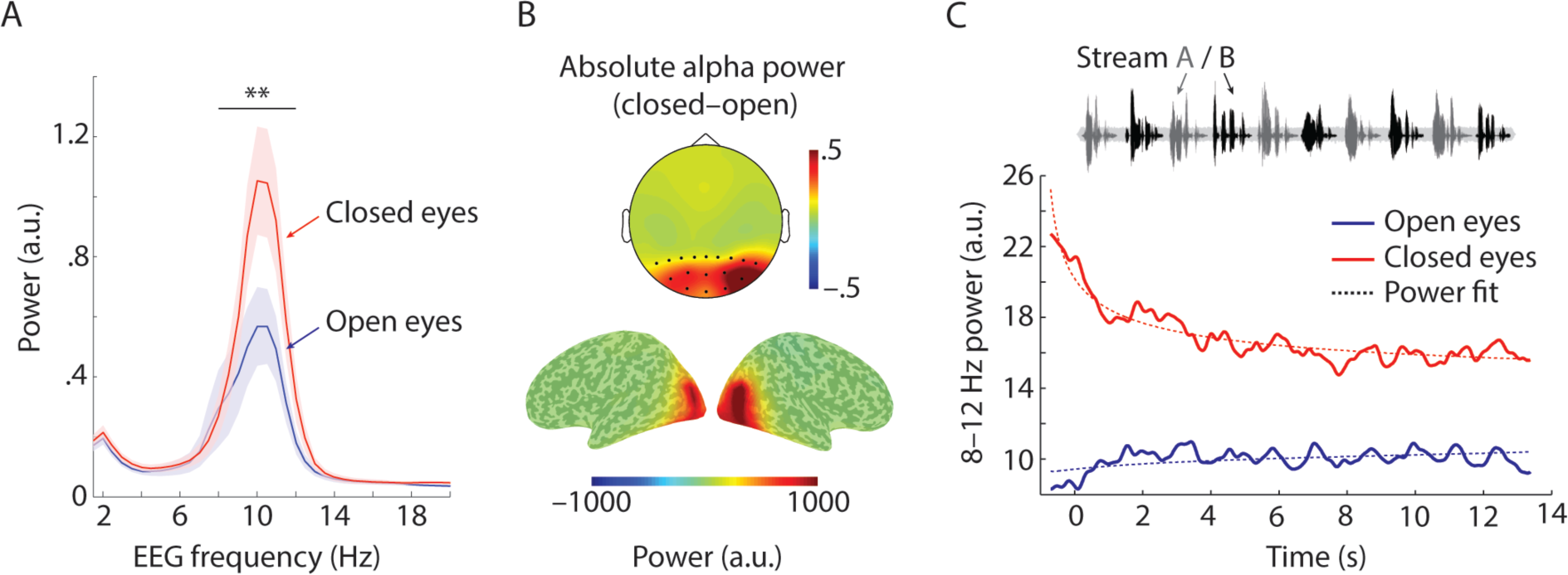
Absolute alpha power in the selective listening task. (A) Average absolute oscillatory power ± 1 between-subject SEM during the trial (0–13.39 s; across *N* = 22 participants and occipital electrodes highlighted in topography in B) increased for closed (red) compared to open eyes (blue) in the alpha frequency band (8–12 Hz; ** *p* = 0.001). (B) Activation maps on inflated brain surfaces show the contrast of 8–12 Hz source-localized power for closed–open eyes. (C) Time courses of oscillatory alpha power averaged across 17 electrodes highlighted in B, separately for eyes-open (blue) and eyes-closed trials (red). Dashed lines show the average across individual participants’ power function fits to the data.

Second, time-resolved power was obtained for four experimental conditions in the 2 (Eyes: open vs. closed) × 2 (Attended stream: A, odd positions vs. B, even position) design. Since we did not expect neural differences of attention to female-vs. male-voiced numbers, we collapsed across the two levels of the factor Attended voice (female vs. male) for all further analyses. Oscillatory power was obtained via calculation of complex Fourier coefficients for a moving time window (fixed length of 0.5 s; Hann taper; moving in steps of 0.02 s through the trial). Power (squared magnitude of complex Fourier coefficients) was obtained for frequencies 1–20 Hz with a frequency resolution of 0.5 Hz.

In order to descriptively characterize time courses of alpha power for open and closed eyes (irrespective of whether participants attended to stream A or B), we fitted a power function (*y = a * (t^m^) + intc*, with t = time, a = scaling parameter, m = exponential coefficient, and intc = intercept) to individual participants’ 8–12 Hz power time courses in the time interval –0.7–13.39 s relative to sound onset (using the *powerfit* function and the least squares procedure implemented in the *lsqcurvefit* function in Matlab).

For all analyses of the attentional modulation of oscillatory power, to remove possible influences of absolute power, single–subject time-frequency representations were averaged across trials and baseline-corrected by subtraction of average power throughout the entire trial (0–13.39 s). To quantify the influence of attention on oscillatory power, we obtained a measure of attentional modulation (Fig. 3B) through subtraction of time-frequency spectra for attention to stream A minus attention to stream B, separately for open versus closed eyes for each participant. To quantify the strength to which attention rhythmically modulated oscillatory power, we calculated Fourier spectra on single subject’s power time-courses (0–13.39 s) obtained from the attentional modulation (Wöstmann et al., 2016). Plotting the amplitudes of Fourier spectra against EEG frequencies revealed modulation spectra (Fig. 3C).

**Figure 3.**
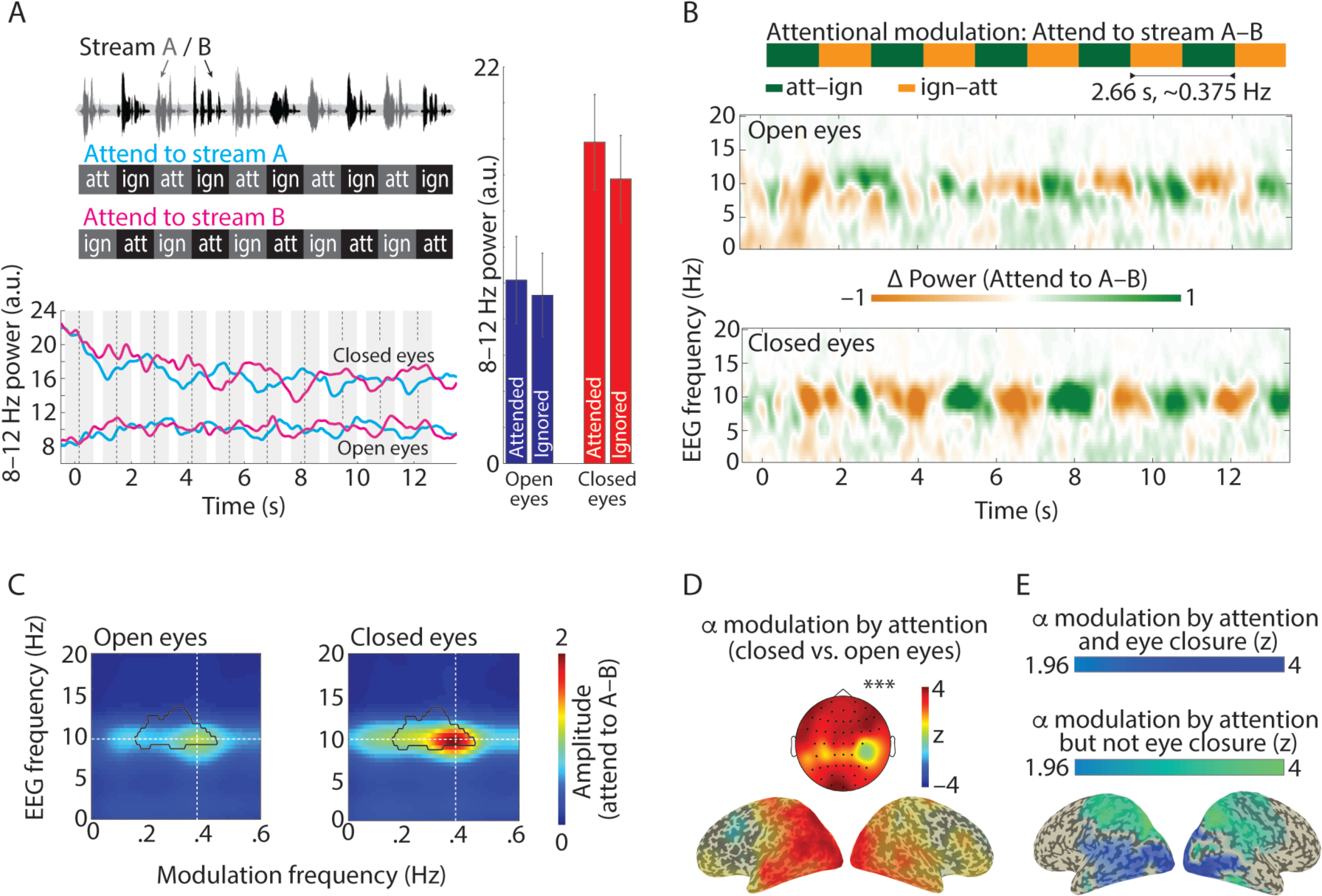
Closing the eyes enhances neural signatures of auditory attention. (A, top) Our task design was intended to induce fluctuationbetweenstates of attending(att) and ignoring (ign) overthe time course of atrial. (A, bottom) Oscillatory alpha power averaged across (*N* = 22) participants and 17 occipito-parietal scalp electrodes (highlighted in Fig. 2B) for trials where participants attended to stream A (cyan) or B (pink) with eyes open (bottom lines) or closed (top lines). Bars and error bars show mean ± 1 between-subject SEM of 8–12 Hz alpha power, averaged across occipito-parietal scalp electrodes and time intervals –0.5 to 0.5s around onsets of attended or ignored numbers (grey shaded areas in line plot). (B) Contrasting trials with attention to stream A–B revealed the attentional modulation, which alternated between states of ‘att–ign’ or ‘ign–att’ at a rate of 0.375 Hz. Time-frequency spectra show the attentional modulation of grand-average oscillatory power (across *N* = 22 participants and 64 scalp electrodes) for open (top) and closed eyes (bottom). (C) Modulation spectra of the attentional modulation show average amplitudes of Fourier spectra calculated on EEG power envelopes for frequencies 1–20 Hz and modulation frequencies 0–0.6 Hz. Strongest power modulation was observed at the intersection of the alpha band (10 Hz) and the attention rhythm (0.375 Hz, dashed lines). Black outlines show a significant positive cluster (*** *p* < 0.001) for closed vs. open eyes, resulting from a cluster permutation test. (D) Cluster *z*-values for EEG and modulation frequency (10 Hz and 0.375 Hz, respectively) are shown on topography (significant electrodes of cluster highlighted) and inflated brain surfaces. (E) Blue colours show voxels that exhibited both, significant absolute alpha power increase for closed compared to open eyes, as well as significant alpha power modulation by attention (see main text for details). Z-values are averaged across effects of closing the eyes and attention on alpha power. Green colours show z-values for the effect of attention on alpha power for voxels that exhibited significant alpha power modulation by attention but not by closing the eyes.

For statistical analysis, single-subject modulation spectra for closed versus open eyes were contrasted using a cluster permutation dependent-samples t-test (Maris & Oostenveld, 2007). This test clusters adjacent bins of modulation spectra and compares the summed t-statistic of the observed cluster against 1000 randomly drawn clusters from the same data with permuted condition labels.

In order to localize neural sources of the hypothesized significant positive cluster, the Linearly Constrained Minimum Variance (LCMV) beamformer approach was used to construct a common filter (based on all trials in all conditions). This common filter was used to project single-trial EEG time courses into source-space. Next, time-frequency spectra and modulation spectra were calculated for each grid point (in the same way as for single electrodes in sensor space; see above). Finally, z-values for the contrast closed versus open eyes were obtained using non-parametric Wilcoxon signed rank tests for EEG frequency 10 Hz and modulation frequency 0.375 Hz for each grid point (Fig. 3D).

### Statistical analyses

We applied parametric *t*-tests when the data conformed to normality assumptions (*p* > 0.05 in Shapiro–Wilk test) and non-parametric Wilcoxon signed rank tests otherwise. For effect sizes, we report *r_equivalent_* (Rosenthal, 1994; Rosenthal & Rubin, 2003), which is bound between 0 and 1. For non-significant results of statistical tests, we additionally report the Bayes Factor (*BF*; obtained in JASP). In brief, a *BF* > 3 gives support for the alternative hypothesis (i.e., the data would be considered 3 times more likely to occur under the alternative hypothesis than under the null), whereas a *BF* < 0.33 gives support for the null hypothesis (Jeffreys, 1939).

## Results

In the main experiment, we presented participants with two alternating streams of five spoken numbers, each. The leading stream (stream A; grey numbers in Fig. 1A) was distinguishable from the lagging stream (stream B; black numbers in Fig. 1A) by talker gender (female vs. male). Every female-voiced number was thus followed by a male-voiced number and vice versa. On each trial, participants were instructed to attend either to the stream spoken by the female or male voice, which was either the leading or lagging stream. Each participant performed 8 blocks (à 20 trials) in a darkened chamber. Participants kept their eyes open or closed during every second block, respectively.

### Closing the eyes shapes dynamics of absolute alpha power

Eye closure affected absolute (i.e., not baseline corrected) alpha power in two ways. First, as expected, 8–12 Hz power across the entire trial time (0–13.39 s) was significantly enhanced for closed compared with open eyes (Fig. 2A; Wilcoxon signed rank test; *z* = 3.26; *p* = 0.001; *r* = 0.7), primarily in occipital cortex regions (Fig. 2B).

Second, and less expected, eye closure also affected the time course of absolute alpha power during a trial (Fig. 2C). For open eyes, alpha power showed the typical moderate increase in the beginning of a listening task (e.g., Henry, Herrmann, Kunke, & Obleser, 2017; Wöstmann, Herrmann, et al., 2015), followed by saturation. For closed eyes, however, alpha power started out much higher and decreased in the beginning of a trial before levelling well above the open-eyes saturation. In order to descriptively trace these alpha power de-/increases, we fitted power functions to individual participants’ alpha time courses. Exponential coefficients of fitted power functions were significantly smaller (i.e., negative instead of positive) for closed compared with open eyes (Wilcoxon signed rank test; *z* = – 2.03; *p* = 0.042; *r* = 0.43), which shows that eye closure reverses the commonly observed alpha power increase at the onset of an attention-demanding listening task into a power decrease.

### Closing the eyes enhances attentional modulation of alpha power

The major hypothesis of this research was not concerned with the absolute level of alpha power but instead with the attentional modulation thereof. As expected, the temporal (0.375-Hz) fluctuation of attending and ignoring speech induced synchronized states of respective high and low alpha power (Fig. 3A) for open and closed eyes. That is, alpha power relatively increased shortly before and initially during the presentation of a to-be-attended number, whereas it relatively decreased later during number presentation and shortly thereafter.

Since to-be-attended and to-be-ignored numbers were placed at opposing positions in trials where participants attended to stream A versus B, the difference (attend to A–B; Fig. 3B) was used to quantify the neural difference in attending versus ignoring, referred to as ‘attentional modulation’ hereafter. The strength of the rhythmic attentional modulation of oscillatory power (i.e., an FFT calculated on the attentional modulation) is shown in modulation spectra in Figure 3C. In line with prior research (Wöstmann et al., 2016), peaks in modulation spectra show that the 10-Hz alpha power envelope reliably tracked the 0.375-Hz temporal progression of attending versus ignoring. Most importantly, a cluster permutation test revealed that the amplitude of rhythmic alpha modulation was stronger with closed compared to open eyes (cluster *p*-value < 0.001). Beamformer source reconstruction localized the significant increase in 10-Hz power modulation at the attention rhythm of 0.375 Hz under closed eyes mainly in occipital and parietal cortex regions (Fig. 3D).

Besides this hypothesized cluster, one additional significant positive cluster was found in the upper alpha band (∼10–12 Hz), however for slow modulation frequencies < 0.05 Hz (cluster *p*-value = 0.01; not shown), which did not correspond with the task-induced rhythmic 0.375 Hz progression of attending and ignoring. No significant negative clusters were found (all cluster *p*-values > 0.15).

It is of note that the attentional modulation of 10-Hz power at 0.375 Hz for closed versus open eyes correlated with the absolute alpha power increase with closed compared to open eyes (*r_Spearman_* = 0.67; *p* < 0.001). This means that participants with stronger increases in absolute alpha power with closed eyes also exhibited stronger increases in the attentional modulation of alpha power for closed compared to open eyes.

### Limited overlap of alpha generators modulated by attention and eye closure

We asked, to what extent alpha generators modulated by closing the eyes would overlap with alpha generators modulated by auditory attention. To this end, z-values for the contrast closed versus open eyes were calculated on alpha power (8–12 Hz) averaged across the entire trial time (0–13.39 s). Neural alpha generators modulated by closing the eyes were defined as voxels with z-values significantly larger than zero (*z* > 1.96; *α* = 0.025; one-sided; uncorrected). Similarly, alpha generators modulated by attention were defined as voxels exhibiting significantly higher 8–12 Hz alpha power modulation at the attentional rhythm of 0.375 Hz compared to the average across EEG frequencies (1–7 and 13–20 Hz) outside the alpha frequency band (*α* = 0.025; one-sided; uncorrected).

Importantly, the intersection of alpha generators modulated by attention and also by eye closure was limited mainly to bilateral occipital cortex regions and inferior parietal and temporal regions in the left hemisphere (blue colours in Fig. 3E). Critically, a large portion of alpha generators in bilateral parietal regions was exclusively modulated by attention but not by closing the eyes (green colours in Fig. 3E).

### Effect of closing the eyes is specific to power, but not phase-locking

As we did not expect any interactive effects of closing the eyes and attention on neural phase, this analysis was entirely exploratory in nature. The rationale was to exclude potential confounds of phase effects on the analysis of oscillatory power.

In addition to the amplitude of the attentional modulation of alpha power, we also analysed the 0.375-Hz phase angles of the 10-Hz power modulation, which did not differ for closed versus open eyes (Parametric Hotelling paired-samples test for equal angular means; *F_40_* = 1.02; *p* = 0.38).

As a control analysis, we tested whether the observed increase in the attentional modulation of neural oscillations with closed eyes was specific to power or whether it would also show up in the phase-locking across trials (which would speak to evoked, rather than induced power modulation; see Wöstmann, Fiedler, & Obleser, 2017). To this end, we performed the very same analysis as shown for power in Figure 3 for inter-trial phase coherence (ITPC; Lachaux, Rodriguez, Martinerie, & Varela, 1999).

During a trial of the selective listening, low-frequency (0–8 Hz) ITPC peaked at the onsets of spoken numbers (Fig. 4A). The attentional modulation of ITPC (Fig. 4B&C) did not show any pattern of rhythmic modulation at the frequency of 0.375 Hz. A cluster permutation test on ITPC modulation spectra did not reveal any clusters of significant differences for the contrast closed versus open eyes (all cluster *p*-values > 0.07).

**Figure 4.**
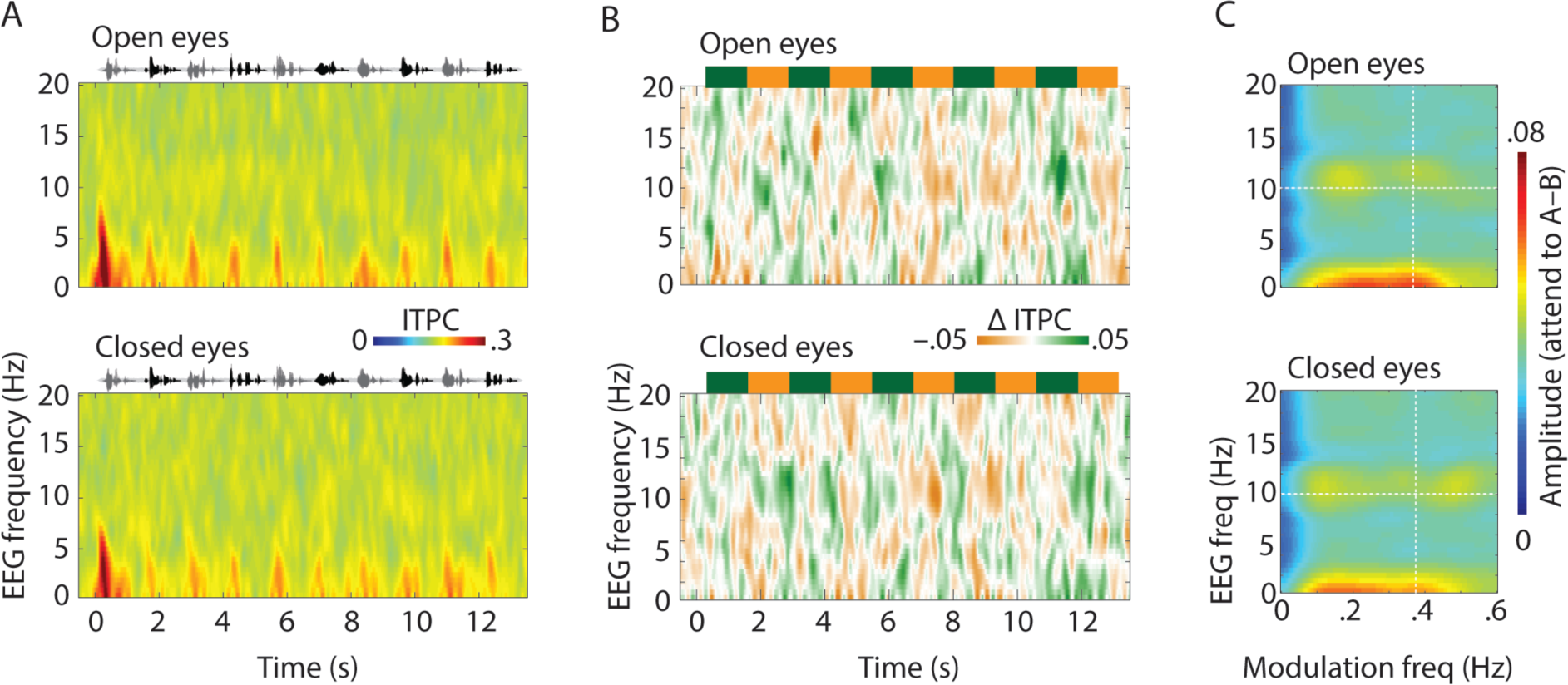
No effect of eye closure on phase-locking of neural oscillations. (A) Average inter-trial phase coherence (ITPC) across *N* = 22 participants and 64 EEG scalp electrodes for trials with eyes open (top) or closed (bottom). (B&C) Same as Fig. 3B&C but for ITPC.

### No sizable relation of attentional alpha power modulation and behaviour

At the end of each trial of the main experiment, we presented participants with three random numbers previously presented in that trial (spoken by a gender-neutral voice). Participants indicated for each of these probe numbers, whether it had been among the to-be-attended numbers (‘Yes, attended’-response) or not (‘No, ignored’-response). We tested whether the attentional modulation of alpha power would relate to behavioural performance.

First, we calculated modulation spectra (see Fig. 3C), separately for correct trials (3 correct responses) and incorrect trials (< 3 correct responses). The attentional modulation of 10-Hz power at 0.375 Hz averaged across 17 occipital electrodes (highlighted in Fig. 2B) was submitted to a 2 (Eyes: open vs. closed) × 2 (Accuracy: correct vs. incorrect) repeated-measures ANOVA. The ANOVA yielded a weak main effect of Accuracy (*F_1, 21_* = 4.17; *p* = 0.054), indicating a tendency for stronger attentional alpha power modulation for correct compared to incorrect trials. The Eyes × Accuracy interaction was not significant (*F_1, 21_* = 0.09; *p* = 0.77). Note, however, that this analysis is not capable of expressing task accuracy in terms of sensitivity and bias, and that results of this analysis might be somewhat confounded by different proportions of correct trials (on average 42%) and incorrect trials (on average 58%).

Thus, in a second analysis, to obtain a more robust within-subject measure of the brain-behaviour relation, we followed a leave-half-out (LHO) subsampling approach: For each participant, we randomly sampled 50% of trials in each condition of the 2 (Attended stream: A vs. B) × 2 (Eyes: open vs. closed) design. We repeated this step 1,000 times to obtain 1,000 LHO estimates of d’ (termed d’_LHO_) and criterion (termed c_LHO_). For the same subsamples we obtained 1,000 LHO estimates of the attentional alpha power modulation (termed attentional alpha modulation_LHO_). Figure 5A shows d’_LHO_ averaged across participants as a function of the rank-ordered attentional alpha modulation_LHO_.

For statistical analysis, we obtained the slope (β) for the linear regression of d’LHO on attentional alpha modulationLHO for each participant. In line with the first analysis, slopes were positive on average (across all conditions: β = 0.02; open eyes: β = 0.025; closed eyes: β = 0.028), suggesting a positive relation of alpha power modulation and perceptual sensitivity (d′). However, slopes were not statistically different from zero (across all conditions: *t_21_* = 1.17; *p* = 0.26; *r* = 0.25; *BF* = 0.41; open eyes: *t_21_* = 1.10; *p* = 0.283; *r* = 0.23; *BF* = 0.38; closed eyes: *t_21_* = 1.18; *p* = 0.25; *r* = 0.25; *BF* = 0.41). Taken together, evidence from these two analyses of the brain–behaviour relation speaks to a weak relation at best of the attentional alpha power modulation at the neural level and task performance at the behavioural level.

Of note, the same analysis procedure did not reveal an obvious relationship of attentional alpha modulationLHO and response bias (c_LHO_; across conditions: *t_21_* = –0.90; *p* = 0.377; *r* = 0.19; *BF* = 0.32; open eyes: *t_21_* = –0.501; *p* = 0.618; *r* = 0.11; *BF* = 0.25; closed eyes: *t_21_* = 1.21; *p* = 0.24; *r* = 0.26; *BF* = 0.43).

### Closing the eyes does not affect behavioural indices of attention and tone detection

Finally, we tested whether closing the eyes would relate to behavioural performance. In the main experiment, perceptual sensitivity in separating to-be-attended from to-be-ignored numbers (d′) was well above zero, i.e., chance level (Fig.5C; open eyes: *t_21_* = 12.19; *p* < 0.001; *r* = 0.94; closed eyes: *t_21_* = 9.36, *p* < 0.001; *r* = 0.9) but did not differ for closed versus open eyes (*t_21_* = 0.45; *p* = 0.654; *r* = 0.1; Bayes Factor, *BF* = 0.24). Participants’ criterion (c) was larger than zero (open eyes: *t_21_* = 1.9; *p* = 0.071; *r* = 0.38; closed eyes: *t_21_* = 2.19; *p* = 0.04; *r* = 0.43), which indicates that participants had an overall conservative bias to respond ‘No, ignored’ more often. However, the criterion was largely unaffected by eye closure (*t_21_* = 1.03; *p* = 0.313; *r* = 0.22; *BF* = 0.36).

We further investigated whether the behavioural null-effects of closing the eyes were specific to a task involving selective listening to speech, or whether they would conceptually replicate in a task that does neither involve speech stimuli, nor the necessity to retain target stimuli in memory.

Accordingly, in the follow-up experiment, participants (*N* = 19) had the task to detect a near-threshold target tone in noise (Fig. 5D). The Signal-to-Noise Ratio (SNR) of the target tone in noise was titrated for each participant before the experiment. The mean titrated SNR was –7 dB (*SD* = 1.55), which resulted in an average proportion correct of 0.646 (*SD* = 0.079) in the experiment.

In line with the main experiment, perceptual sensitivity (d’) in the follow-up experiment was significantly above zero (Fig. 5E&F; open eyes: *t_18_* = 7.57; *p* < 0.001; *r* = 0.87; closed eyes: *t_18_* = 7.42; *p* < 0.001; *r* = 0.87) but did not differ for open compared with closed eyes (*t_18_* = 0.899; *p* = 0.381; *r* = 0.207; *BF* = 0.339). The response criterion (c) was significantly positive (open eyes: *t_18_* = 10.32; *p* < 0.001; *r* = 0.92; closed eyes: *t_18_* = 8.03; *p* < 0.001; *r* = 0.88), indicating a generally conservative bias to respond ‘No’ (no tone detected) more often. The response criterion did not differ for open versus closed eyes (*t_18_* = 1.083; *p* = 0.293; *r* = 0.247; *BF* = 0.397).

In addition to objective measures of behavioural task performance, we assessed participants’ subjective experience of the listening tasks with closed versus open eyes. Such subjective ratings are of interest, as we have recently shown that higher (prestimulus) alpha power correlates with lower subjective ratings of confidence in a pitch discrimination task (Wöstmann, Waschke, & Obleser, 2018). After each block of the main experiment, participants rated their estimated task performance, effort, and tiredness on a scale from 1 to 10. Average ratings did not differ significantly for blocks with closed versus open eyes (estimated task performance: *t_21_* = 0.11; *p* = 0.914; *r* = 0.02; *BF* = 0.224; effort: *z* = 0.23; *p* = 0.821; *r* = 0.12; *BF* = 0.258; tiredness: *t_21_* = 1.23; *p* = 0.234; *r* = 0.26; *BF* = 0.434). After the follow-up experiment, participants rated the task ease on a scale from 1 to 6 (1 = easier with closed eyes; 6 = easier with open eyes). Ratings did not differ significantly from 3.5, that is, the theoretical centre of the scale (*z* = 1.21; *p* = 0.225; *r* = 0.23; *BF* = 0.373).

The small Bayes Factors (*BFs*) for the non-significant comparisons of sensitivity, criterion, and subjective ratings for closed versus open eyes in the main experiment as well as in the follow-up experiment lend support to the null hypothesis of no difference in objectively assessed as well as subjectively rated behavioural performance in auditory attention and tone detection with closed versus open eyes.

## Discussion

The present study sought to test whether closing the eyes during listening has the potency to enhance neural and behavioural indices of auditory attention. The major findings can be summarized as follows. As expected, closing the eyes during listening increased the level of absolute alpha power. Importantly, however, eye closure amplified the neural alpha power difference in attending versus ignoring speech, that is, the neural separation of relevant and irrelevant acoustic input. Finally, debunking the belief that eye closure per se improves listening, we provide evidence for the absence of an effect of closing the eyes on sensitivity and response criterion for auditory attention and tone detection in two independent experiments.

### Closing the eyes shapes absolute alpha power dynamics

In agreement with previous research (e.g., Barry, Clarke, Johnstone, Magee, & Rushby, 2007; Geller et al., 2014), eye closure in a darkened room (Adrian & Matthews, 1934; Ben-Simon et al., 2013) almost doubled participants’ absolute occipital alpha power in the EEG (Fig. 2). This absolute alpha power increase possibly reflects inhibition of neural processing in visual cortex regions (Foxe & Snyder, 2011; Jensen & Mazaheri, 2010), disengaging these regions when the eyes are closed. Of note, Adrian (1944) was possibly the first to discover that alpha oscillations show a general increase when participants direct their attention to the auditory modality (for more recent studies, see e.g., Dimitrijevic, Smith, Kadis, & Moore, 2017; Foxe, Simpson, & Ahlfors, 1998; Henry et al., 2017; Wöstmann, Herrmann, et al., 2015). He interpreted this alpha increase as “a positive activity that fills those parts of the cortex which are for the moment unemployed.”^2^

We here observe that eye closure propels alpha power outside the dynamic range observed with open eyes (Fig. 2C). Since we used eye closure in this study as a means to modulate alpha power, it can be considered a method of endogenous neuromodulation, comparable to (exogenous) perturbation methods such as transcranial alternating current stimulation (tACS; Herrmann, Rach, Neuling, & Struber, 2013). However, eye closure appears to induce much stronger increases in alpha power, which was also demonstrated by Neuling et al. (2013), who showed that alpha-tACS increases neural alpha power only in a regime of low alpha power with open eyes but not in a regime of high alpha power with closed eyes.

In sum, eye closure lifted absolute alpha power far above levels observed during attentive listening with open eyes.

### Closing the eyes enhances the attention-induced alpha modulation

In the present selective listening task (main experiment), a listener’s intent to attend versus ignore spoken numbers was accompanied by respective states of high versus low alpha power (Fig. 3). This agrees with previous findings of alpha power modulation in temporal synchrony with attending versus ignoring speech (Tune, Wöstmann, & Obleser, 2018; Wöstmann et al., 2016). Our results speak to the modulation of alpha power over time as a neural signature of auditory attention.

The most important finding of this study was that eye closure not only increased the overall level of absolute alpha power, but also strongly enhanced the attentional modulation thereof. Enhanced alpha power modulation with closed eyes was generally wide-spread in topography but localized mainly in non-auditory, parieto-occipital cortex regions. Alpha power increases in these regions have been associated with inhibitory control of supramodal attention networks (e.g., Banerjee, Snyder, Molholm, & Foxe, 2011) and visuo–spatial processing areas (e.g., Fu et al., 2001; Wöstmann, Herrmann, et al., 2015). In visual tasks, alpha power modulation in parieto-occipital regions in sync with the stimulus affects stimulus encoding (Bonnefond & Jensen, 2012; Park et al., 2014; Payne, Guillory, & Sekuler, 2013). In the present auditory task, however, parieto-occipital alpha power modulation might control the degree of auditory attention through inhibition of task-irrelevant visual processing areas (Strauß et al., 2014). The present results show that closing the eyes is indeed effective insofar as it enhances the degree to which a listener’s top-down intention to attend versus ignore speech modulates neural alpha power.

The effect of closing the eyes was specific to the 8–12 Hz alpha frequency range, and observed only for power but not for phase-locking of neural oscillations. This speaks against the possibility that neural responses in general are enhanced with closed compared to open eyes. Furthermore, this indicates that cycles of the eye–closure-modulated alpha oscillation were not evoked (i.e., strictly phase-locked across trials) but rather induced (for distinction of evoked and induced oscillations, see Tallon-Baudry & Bertrand, 1999; Wöstmann et al., 2017).

So, what is the functional significance of larger alpha power modulation with closed eyes? It has been suggested that even in a darkened room with very little visual input, closing the eyes might reduce the cross-modal dominance of the visual system (Brodoehl, Klingner, & Witte, 2015) to facilitate attention to non-visual modalities. Eye closure could thus free neural processing capacity, in order to enhance inhibition versus facilitation of respective ignored versus attended auditory stimuli, which eventually surfaces in stronger alpha power modulation. Neural separation of attended and ignored auditory stimuli is a key mechanism of auditory attention (Shinn-Cunningham, 2008) to solve the long-standing Cocktail party problem, that is, listening to one talker against competing distractors (Cherry, 1953). In this respect, alpha power modulation has been considered a filter mechanism to separate attended from ignored auditory input (Kerlin, Shahin, & Miller, 2010; Strauß et al., 2014; Wöstmann et al., 2016). The present results demonstrate that this auditory filter mechanisms surfaces to a stronger extent in the human EEG signal when the eyes are closed.

### Effects of eye closure on neural but not behavioural responses

Against the widely held belief that eye closure improves attentive listening, we found no increase in listeners’ sensitivity or criterion in telling attended from ignored speech items in the main experiment (Fig. 5). In principle, these null effects might have been caused by nature of our task. Spoken numbers were presented well above participants’ hearing thresholds and sensitivity of responses by far exceeded chance level. This contrasts with previous studies in the somatosensory modality where eye closure was found to lower perceptual detection at threshold (Brodoehl, Klingner, Stieglitz, et al., 2015; Brodoehl, Klingner, & Witte, 2015).

Furthermore, it might be that potential effects of closing the eyes on auditory processing were masked by the fact that task performance in the main experiment depends to some extent also on working memory for the presented numbers. However, the follow-up experiment, which implemented a close-to-threshold tone detection task and was free of potential influences of working memory, conceptually replicated the behavioural null-effects of the main experiment.

Post-hoc correlation analyses revealed that 14–18% of variance in sensitivity (d-prime) in the main experiment were explained by established tests of working memory capacity (i.e., backward and forward digit span scores, respectively). Statistical control for participants’ working memory capacity had virtually no influence on the significant effect of eye closure on alpha power modulation, as well as on the non-significant effect of eye closure on behaviour. In line with the general view that attention and working memory interact strongly (Awh, Vogel, & Oh, 2006), we argue that both of these processes serve to focus on task-relevant numbers and to suppress distractors in the present task. Since target and distractor numbers were separated in time, the major demand on attention was not to segregate numbers in the auditory periphery. Rather, the demand on attention was to selectively store target numbers in working memory and to protect working memory content against distractor interference. Therefore, although the task in the main experiment has a clear working memory component, it critically depends on attention and can thus be considered an auditory selective attention task.

We consider three potential reasons for why eye closure was found here to affect neural but not behavioural indices of auditory attention and perception: Issues with sensitivity of our behavioural outcome measure; a mismatch of the imposed neuromodulation (eye closure) and the required neural dynamic range (Jazayeri & Afraz, 2017); and the absence of visual distraction.

First, it might be that our behavioural outcome measures were not sensitive enough to detect any effect of eye closure. We consider this possibility rather unlikely since (i) we found the null-effect both in an auditory selective listening task (main experiment), as well as in a tone detection task (follow-up experiment), (ii) our analyses of sensitivity and response criterion had the power to detect changes in behavioural performance that might be overseen in case only the proportion of correct responses is inspected, and (iii) our Bayesian statistical analysis revealed small Bayes Factors (close to or below .33; Dienes, 2014) for behavioural null-effects, which provide positive evidence for the absence of effects rather than insensitivity of the data. Note, however, that our behavioural null-results do not preclude the possibility that eye closure might be beneficial in other listening scenarios, such as tasks involving auditory spatial attention.

Second, we observed only a weakly positive association between the attentional modulation of alpha power and perceptual sensitivity (Fig. 5A). Thus, increases in the attentional modulation of alpha power do not directly result in sizable improvements of behavioural performance. Furthermore, our results suggest that occipital alpha power generators that increased their modulation by closing the eyes did not precisely overlap with parietal alpha generators associated with auditory attention (Fig. 3E). It is known that the impact of neuromodulation on behaviour critically depends on its precision in targeting task-relevant neural processes in time, frequency, and space (e.g., Herrmann, Murray, Ionta, Hutt, & Lefebvre, 2016; Vosskuhl, Struber, & Herrmann, 2018). Furthermore, neuromodulation might be behaviourally ineffective if it pushes the neurally relevant process outside its natural dynamic range (Jazayeri & Afraz, 2017). In the present study, it might be that eye-closure did not affect behavioural performance since the eye-closure induced effect on alpha oscillations was limited to occipital regions and thus spatially distinct from parietal alpha generators involved in auditory attention.

Third, it might be the case that our experimental intervention of closing the eyes was indeed successful in increasing versus decreasing the inhibition of non-auditory regions during respective periods of attending versus ignoring speech. However, our participants performed the main- and follow-up experiment in a darkened chamber, deprived of visual input. Thus, inhibition of non-auditory regions might have been ineffective in modulating behaviour since no non-auditory, distracting stimuli were presented. Although the intention of the present study was to study the sole effect of closing the eyes free from potential influences of the blocking of the visual input, future studies might observe beneficial effects of closing the eyes on listening tasks in more busy visual environments.

## Conclusions

Does closing the eyes enhance auditory attention? First, an important neural mechanism of auditory attention, alpha power modulation, has been shown here to be amplified by eye closure. That is, when listeners have their eyes closed, the increased attentional modulation of alpha power indeed implies stronger neural separation of attended and ignored sound sources. Second, however, eye closure per se does not improve auditory attention and tone detection performance. Possibly, the impact of eye closure on neural oscillatory dynamics does not match alpha power modulations associated with listening performance precisely enough. Third, the present findings do have important practical implications for the neuroscience of auditory attention: Researchers should rigorously control whether participants close their eyes during listening. Participants might even be instructed to utilize the endogenous amplification of closing the eyes to increase the power to observe existing attentional effects on neural alpha oscillations.

## Footnotes

1 Cherry CE (1953) Some Experiments on the Recognition of Speech, with One and with Two Ears. Journal of the Acoustical Society of America 25:975-979. P. 976

2 (Adrian, 1944) P. 361

